# Impeding transcription of expanded microsatellite repeats by deactivated Cas9

**DOI:** 10.1101/185496

**Authors:** Belinda S. Pinto, Tanvi Saxena, Ruan Oliveira, Héctor R. Méndez-Gómez, John D. Cleary, Lance T. Denes, Ona McConnell, Juan Arboleda, Guangbin Xia, Maurice S. Swanson, Eric T. Wang

## Abstract

Transcription of expanded microsatellite repeats is associated with multiple human diseases, including myotonic dystrophy, Fuchs’ endothelial corneal dystrophy, and *C9orf72*-ALS/FTD. Eliminating or reducing production of RNA and proteins arising from these expanded loci holds therapeutic benefit. Here, we tested the hypothesis that a deactivated form of the Cas9 enzyme impedes transcription across expanded microsatellites. We observed a repeat length-, PAM-, and strand-dependent reduction in the abundance of repeat-containing RNAs upon targeting dCas9 directly to repeat sequences. Aberrant splicing patterns were rescued in DM1 cells, and production of RAN peptides characteristic of DM1, DM2, and *C9orf72*-ALS/FTD cells was drastically decreased. Systemic delivery of dCas9/gRNA by adeno-associated virus led to reductions in pathological RNA foci, rescue of chloride channel 1 protein expression, and decreased myotonia. These observations suggest that transcription of microsatellite repeat-containing RNAs is more sensitive to perturbation than transcription of other RNAs, indicating potentially viable strategies for therapeutic intervention.

## Introduction

Microsatellite expansion diseases are a class of genetically inherited conditions associated with the destabilization and expansion of short repetitive sequence elements in the genome, which cause pathogenic effects via multiple mechanisms including epigenetic silencing, RNA gain of function, and/or protein gain of function^1^. These diseases, which include myotonic dystrophy Types 1 and 2 (DM1, DM2), Fuchs’ endothelial corneal dystrophy (FECD), Huntington Disease (HD), *C9orf72*-ALS/FTD (C9ALS/FTD) and spinocerebellar ataxias (SCAs), are often multi-systemic and can profoundly affect the central nervous system, muscle, and the heart. Somatic instability causes repeat expansion throughout the lifetime of an individual, with the most dramatic expansions reaching thousands of nucleotides in length in those post-mitotic tissues^2^,^3^. Cellular toxicity in these diseases partly occurs due to transcription of the expanded repeat tract. For example, in DM1 and DM2 expanded CUG or CCUG repeat RNAs, respectively, sequester Muscleblind-like (MBNL) proteins away from their endogenous RNA targets^4^ leading to aberrant splicing patterns^5^ and altered RNA stability/localization^6^,^7^, among other effects. In C9ALS/FTD, expanded G_4_C_2_ and C_4_G_2_ repeat RNAs sequester several RNA binding proteins, as well as undergo RAN translation to produce dipeptide polymers toxic to cells^8^^-^^10^. Taken together, these findings suggest that silencing of the expanded repeat locus holds therapeutic value.

Various approaches have been taken to silence toxic RNA or protein in microsatellite expansion diseases, including antisense oligonucleotides^5^,^11^^-^^13^, small RNAs^14^^-^^17^, and small molecules^18^,^19^. Perturbation to co-factors of RNA polymerase II (RNA Pol II) reduces transcription through expanded repeats in HD^20^ and C9ALS/FTD^21^ models, and treatment with Actinomycin D at nanomolar doses preferentially impedes transcription of CTG repeats in DM models^18^. A prevailing hypothesis is that the efficiency of transcription through expanded repeats is decreased relative to non-repetitive sequences. This would provide a therapeutic window through which to impede transcription of these sequences in a repeat length-dependent manner resulting in premature termination and nascent transcript turnover.

A deactivated version of the Cas9 enzyme, of the clustered regularly interspaced short palindromic repeats (CRISPR) system, has been used to impair transcription of specific loci, as well as to visualize, tether, and/or isolate DNA in a sequence-specific manner. In prokaryotes, deactivated Cas9 (dCas9) can efficiently inhibit transcriptional initiation and elongation when bound to gene bodies or promoters^22^. In eukaryotes, dCas9 can inhibit transcriptional initiation when fused to an inhibitory domain and targeted near the transcription initiation site^23^^-^^26^. However, elongation inhibition by targeting dCas9 alone to the gene body has been largely ineffective, even when recruiting dCas9 to > 90 possible targeting sites^27^.

Here, we test the hypothesis that expanded microsatellite repeats are highly sensitive to transcriptional blockade by dCas9, even in the context of processively elongating RNA Pol II. We find that the efficiency of inhibition follows rules similar to that observed in non-repetitive contexts, with clear dependencies on proto-spacer adjacent motif (PAM) sequence and the targeted DNA strand. In addition, application of this approach to cell and animal models of disease rescues downstream pathogenic consequences. These observations present a novel application of the CRISPR/Cas9 system, provide new tools to impede transcription of expanded microsatellites in a strand-dependent manner, and suggest that targeting transcription in a repeat length-dependent manner may be a viable therapeutic strategy for these diseases.

## Results

### dCas9-gRNA complexes reduce abundance of repeat-containing RNAs in a length-, PAM-, and strand-dependent manner

Binding of deactivated *S. pyogenes* Cas9 (dCas9) to DNA in the vicinity of transcriptional start sites (TSSs) can inhibit transcription in prokaryotes and eukaryotes when coupled with a transcriptional inhibitory domain. However, inhibition of transcriptional elongation via gene body targeting (>1kb away from TSS) has been relatively inefficient in eukaryotes. We hypothesized that transcriptional inhibition of expanded microsatellite repeats by dCas9 would be more potent as the repetitive nature of these repeats, sometimes thousands of tandem copies, would: 1) present challenges for RNA Pol II elongation even under normal conditions; and 2) allow for high levels of dCas9 recruitment using a single guide RNA (gRNA) sequence forming a substantial block to the elongating polymerase (**Fig. 1A**). To test this model, we employed a plasmid-based strategy in cell culture to examine the effects of dCas9 recruitment on transcription of CTG/CAG repeats that occur in DM1, FECD, HD, and SCAs. We utilized plasmids containing 0, 12, 40, 240, 480, or 960 CTG repeats located >1.5 kilobases downstream of the TSS in the *DMPK 3’* UTR. To recruit dCas9 to these repeats, we designed gRNAs targeting CTG/CAG repeats in each of 3 possible nucleotide phases of the non-template strand, and 1 phase of the template strand (**Fig 1B**, **left panel**). Each CRISPR PAM was, therefore, constrained to CAG, AGC, GCA, or CTG. To precisely measure the expression of the repeat transcripts, we developed an amplicon-based deep sequencing assay called “<u>M</u>easurement of <u>B</u>arcoded <u>T</u>ranscripts by <u>A</u>mplicon Sequencing” (MBTA-Seq) (**Fig. S1**). For this assay, we modified the plasmids by introducing distinct 8 nucleotide barcodes downstream of each CTG repeat tract. RNA was harvested from cells transfected with these plasmids, polyA+ selected, amplified by RT-PCR across the barcode region but avoiding the repeat containing region, and deep sequenced (**Fig. S1**). This assay was highly reproducible, and allowed us to simultaneously measure the expression of multiple repeat-containing transcripts in the same pool of cells (**Table S1**). In the presence of dCas9 and (CAG)_6_ gRNA, we observed a dramatic reduction in expression of RNAs containing expanded CUG repeats (**Fig. 1B**, **right panel**). Knockdown efficiency increased as a function of the number of repeats present, presumably as a function of the number of dCas9-gRNA complexes that could be recruited. Twelve CTG repeats, likely recruiting at most a single dCas9-gRNA complex, showed ~50% repeat-containing RNA remaining, and 40 CTG repeats showed ~25% remaining. Repeat lengths >240 CTG showed only ~5% remaining. (AGC)_6_ and (GCA)_6_ gRNAs showed poor knockdown efficiency, consistent with previously described SpCas9 PAM preferences, where NGG is best, NAG is second best, and NCG/NTG are equally disfavored^28^. The (CUG)_6_ gRNA showed little to no knockdown of transcripts containing expanded CUG repeats, potentially due to a weak PAM as well as targeting to the template strand. Importantly, dCas9 was required for decreased expression of repeat-containing transcripts, as presence of gRNAs alone did not lead to knockdown (**Fig. S2A**).

**Figure 1.**
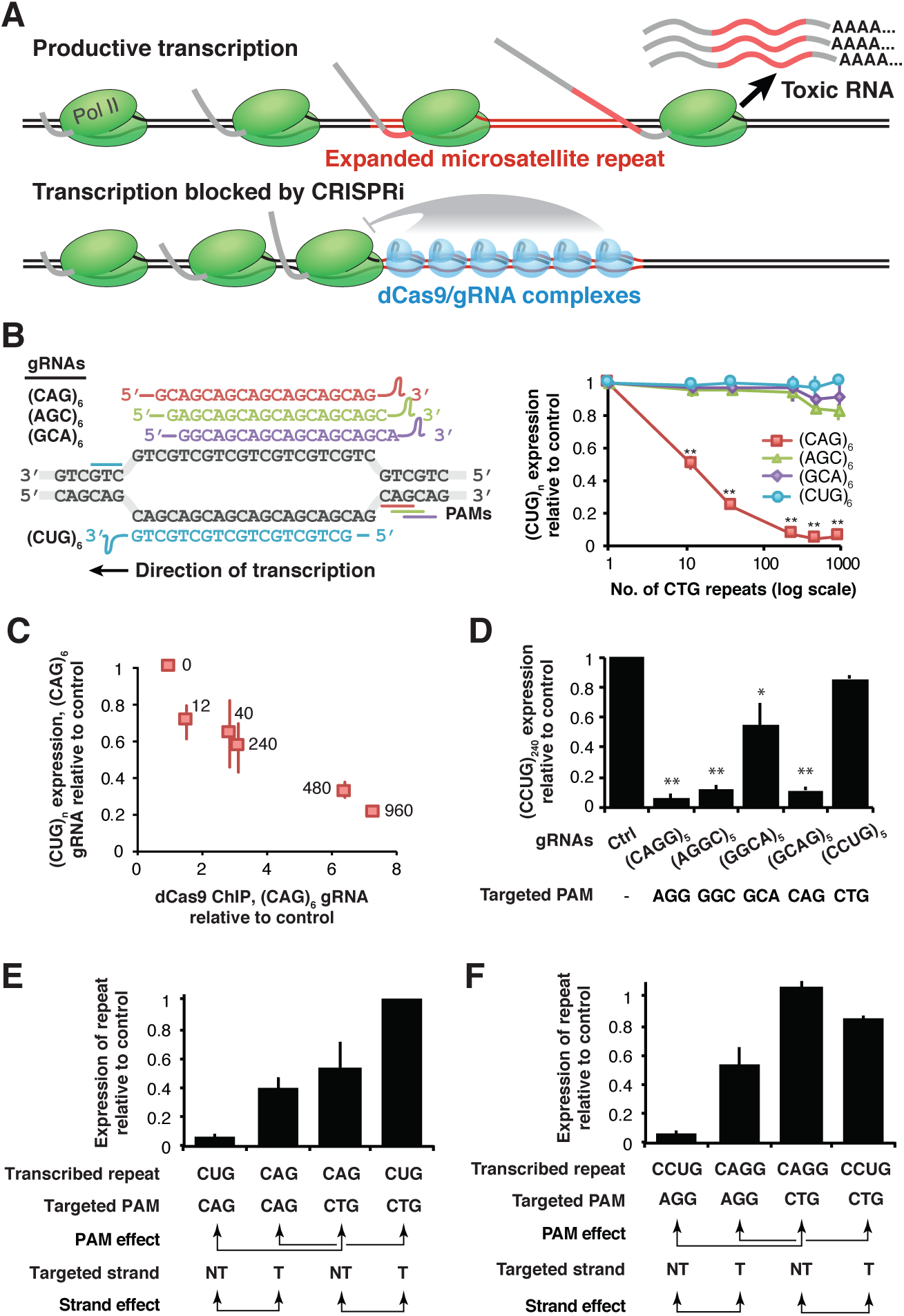
Deactivated SpCas9 impedes transcription of expanded microsatellite repeats in a length-, PAM-, and strand-dependent manner. A) Proposed model for how recruitment of multiple dCas9/gRNA complexes to expanded microsatellite repeats impedes transcription by RNA Pol II B) Schematic of gRNAs used to target transcription of CTG repeats (left panel). Abundance of CUG repeat-containing RNA in the presence of dCas9/gRNAs targeting the repeat tracts of various lengths in HeLa cells, relative to the same RNA species with zero repeats (right panel). Error bars show standard deviation. C) Relative dCas9 ChIP signal across all repeat lengths versus percent RNA remaining following (CAG)_6_ gRNA treatment relative to control gRNA treatment. dCas9 ChIP signal is computed as dCas9 immunoprecipitation divided by input chromatin in the presence of (CAG)_6_ gRNA, divided by dCas9 immunoprecipitation divided by input chromatin in the presence of control gRNA. Relative abundance of repeat-containing loci and RNAs was assessed by deep sequencing of associated barcodes. Error bars show standard error of the mean. D) Abundance of CCUG repeat-containing RNA in the presence of dCas9 and various gRNAs targeting the (CCTG)_240_ repeat tract in HeLa cells, relative to the same RNA species with zero repeats. E) Abundance of RNAs containing 960 CUG repeats or 960 CAG repeats in the presence of dCas9 and (CUG)_6_ or (CAG)_6_ gRNAs in HeLa cells, relative to RNA species with zero repeats. Arrows denote comparisons relevant for assessing PAM-or strand-dependent effects on efficacy. F) Abundance of RNAs containing 240 CCUG repeats or 240 CAGG repeats in the presence of dCas9 and (CCUG)_5_ or (CAGG)_5_ gRNAs in HeLa cells, relative to RNA species with zero repeats. Arrows denote comparisons relevant for assessing PAM-or strand-dependent effects on efficacy. (two-tailed T-test, **p < 0.0005, *p < 0.005)

These observations suggest that recruitment of dCas9-gRNA complexes can impede transcription of expanded microsatellite repeat tracts in a repeat length-dependent manner. However, these experiments were performed using transiently transfected plasmids, which may not accurately model all aspects of transcriptional regulation in a genomic context. Furthermore, experiments to assess binding of protein complexes to DNA loci are most commonly performed using genomic targets and not plasmids. Therefore, we established a HeLa cell line in which transgenes for each of the 6 CTG repeat lengths, with associated barcodes, were stably integrated (**Fig. S2B**). To measure DNA binding by dCas9-gRNA complexes, we performed chromatin immunoprecipitation (ChIP) against dCas9 in the presence of (CAG)_6_ gRNA or non-targeting control gRNA. We used deep sequencing of barcodes, similar to MBTA-Seq, to quantitate abundance of immunoprecipitated DNA encoding each repeat length. To also confirm transcriptional repression in this cell line, we measured RNA abundance by MBTA-Seq following transfection of dCas9-gRNA complexes. We observed that relative dCas9-gRNA binding to the DNA increased as a function of repeat length, concomitant with a decrease in relative RNA abundance (**Fig. 1C**). In this genomic context, dCas9-gRNA reduced the abundance of transcripts containing 960 CUG repeats to ~20%, and was associated with ~8-fold increased binding relative to genomic loci with no CTG repeat tracts. These observations suggest that binding of dCas9-gRNA complexes to DNA impedes transcription of repeat tracts, and that both binding and potency of transcriptional inhibition increases with the number of repeats.

To test the ability of dCas9-gRNA complexes to impede transcription of other repeat-containing sequences, we assayed CCTG repeat tracts, which cause DM2. dCas9-gRNA complexes similarly reduced expression of CCUG repeat-containing RNAs (**Fig. 1D**). Here, the tetranucleotide repeat allowed testing of four distinct gRNAs targeting the non-template strand, (CAGG)_5_, (AGGC)_5_, (GGCA)_5_, and (GCAG)_5_, corresponding to PAMs AGG, GGC, GCA, and CAG, respectively. Consistent with PAM preferences of SpCas9, knockdown was most efficient with the (CAGG)_5_ gRNA, although the (AGGC)_5_ and (GCAG)_5_ gRNAs were also quite effective. The (CCUG)_5_ gRNA was not effective at impeding transcription of CCTG repeats. Similar to our studies of expanded CUG repeats, presence of gRNAs alone did not lead to knockdown (**Fig. S2C**).

In both bacterial and mammalian systems, blockade of RNA Pol II has been shown to be most effective when targeting dCas9 to the non-template strand of transcribed DNA, likely because helicase activity native to RNA Pol II can unwind nucleic acids hybridized to the template strand^22^. Our results from targeting repeats with (CAG)_6_ gRNA versus (CUG)_6_ gRNA and (CAGG)_6_ gRNA versus (CCUG)_6_ gRNA are consistent with this model, but these comparisons involve changes to both the targeted strand as well as the PAM. In addition, some of the most efficient gRNAs are those that could also facilitate dCas9 targeting of RNA via complementarity between the gRNA and the repeat-containing RNA. To study how the PAM sequence and targeted strand influence the efficacy of transcriptional blockade in isolation, and to clarify DNA versus RNA-targeting mechanisms, we generated (CAG)_960_ and (CAGG)_240_ plasmids suitable for MBTA-Seq. These constructs are identical to their (CTG)_960_ and (CCTG)_240_ repeat-containing counterparts except for their repeat tract and barcode sequences. We then used MBTA-Seq to measure the efficiency of CUG_960_ or CAG_960_ knockdown in the presence of either (CAG)_6_ or (CUG)_6_ gRNA (**Fig. 1E**). By testing all four combinations, we could separate PAM-dependent effects from strand-dependent effects. Consistent with previous reports, targeting the non-template DNA strand reduced expression more effectively than targeting the template strand. Specifically, (CUG)_960_ was reduced to ~5% by a (CAG)_6_ gRNA, while (CAG)_960_ was reduced to ~40% by the same gRNA. These two conditions effectively compare targeting of non-template vs. template strand, controlling for the CAG PAM. Similarly, (CAG)_960_ was reduced to ~50% by the (CUG)_6_ gRNA, and (CUG)_960_ remained relatively high at ~97%. Again, targeting of the non-template strand is more efficient than targeting of the template strand, controlling for the CTG PAM.

PAM-dependent effects were quantified by comparing abundance of (CTG)_960_ in the presence of (CAG)_6_ gRNA relative to abundance of (CAG)_960_ in the presence of (CUG)_6_ gRNA. Here, both gRNAs target the non-template strand, but use different PAMs. We observed ~5% RNA remaining with the CAG PAM, and ~55% RNA remaining with the CTG PAM, controlling for targeted strand, consistent with the CAG PAM being more effective. Similarly, measurement of (CAG)_960_ RNA in the presence of (CAG)_6_ gRNA and (CTG)_960_ RNA in the presence of (CUG)_6_ gRNA allows comparison of both PAMs, controlling for targeted strand. Again, we observed more effective silencing with a CAG PAM (~40% RNA remaining) as compared to the CTG PAM (no change in RNA). We observed similar trends with CCTG/CAGG repeats (**Fig. 1F**). (CCUG)_240_ was reduced to ~5% by (CAGG)_5_ gRNA and (CAGG)_240_ to ~55% by (CAGG)_5_ gRNA (non-template vs. template strand, AGG PAM). We did not observe the same template vs. non-template effect when silencing with a CTG PAM, but the (CCTG)_5_ gRNA was largely ineffective, showing little to no silencing.

Overall, these results separate the effects of PAM sequence and targeted strand in the context of transcriptional blockade. Furthermore, they support a model in which dCas9/gRNA complexes target repeat-containing DNA, because reductions in RNA abundance are achieved even when using gRNAs that are not complementary to transcribed RNAs.

### dCas9-mediated transcriptional inhibition rescues splicing defects and blocks RAN translation in cell-based models of DM and C9ALS/FTD

Downstream symptoms of many repeat expansion diseases are caused by the expression of toxic RNA species. In some cases, such as DM and ALS/FTD, the RNA sequesters RNA binding proteins necessary for cellular functions, leading to numerous downstream molecular changes to the transcriptome and proteome^1^,^4^^-^^10^. To assess the extent to which dCas9-mediated transcriptional silencing can rescue molecular and cellular phenotypes in disease models, we used a HeLa cell-based model of DM1, in which (CUG)_480_ repeats are expressed from a plasmid^29^. Co-expression of dCas9 and (CAG)_6_ gRNA led to a reduction in the percentage of cells showing CUG-containing RNA foci, as well as a reduction in the number of foci per cell (**Fig. 2A, B**). We measured the extent to which MBNL-dependent splicing misregulation, characteristic of DM1 cells^30^^-^^32^, could be rescued using a splicing minigene reporter containing MBNL1 exon 5^33^. The percent spliced in (psi, Ψ) of this exon is regulated by MBNL proteins, and changes from 10% in healthy HeLa cells to ~70% in cells expressing (CUG)_480_ repeats. Co-transfection of dCas9 and (CAG)_6_ gRNA partially rescued splicing dysregulation, reducing Ψ to ~35% (**Fig. 2C**). This rescue occurred only in the presence of both dCas9 and (CAG)_6_ gRNA, indicating that (CAG)_6_ gRNA expression alone cannot neutralize toxic CUG RNA by directly hybridizing to repeat RNA and displacing MBNL protein or facilitating degradation by dsRNA-sensitive nucleases. It is well established that splicing defects in DM1 depend on the extent of MBNL sequestration and therefore total CUG repeat load; Ψ values for MBNL1 exon 5 increased with the length of the CTG repeat tract transfected into cells (Fig. 2D) ^34^. Interestingly, co-expression of dCas9 and (CAG)_6_ gRNA yielded a similar rescue of splicing across all repeat lengths exhibiting mis-splicing, i.e. 240, 480, and 960 CUG repeats.

**Figure 2.**
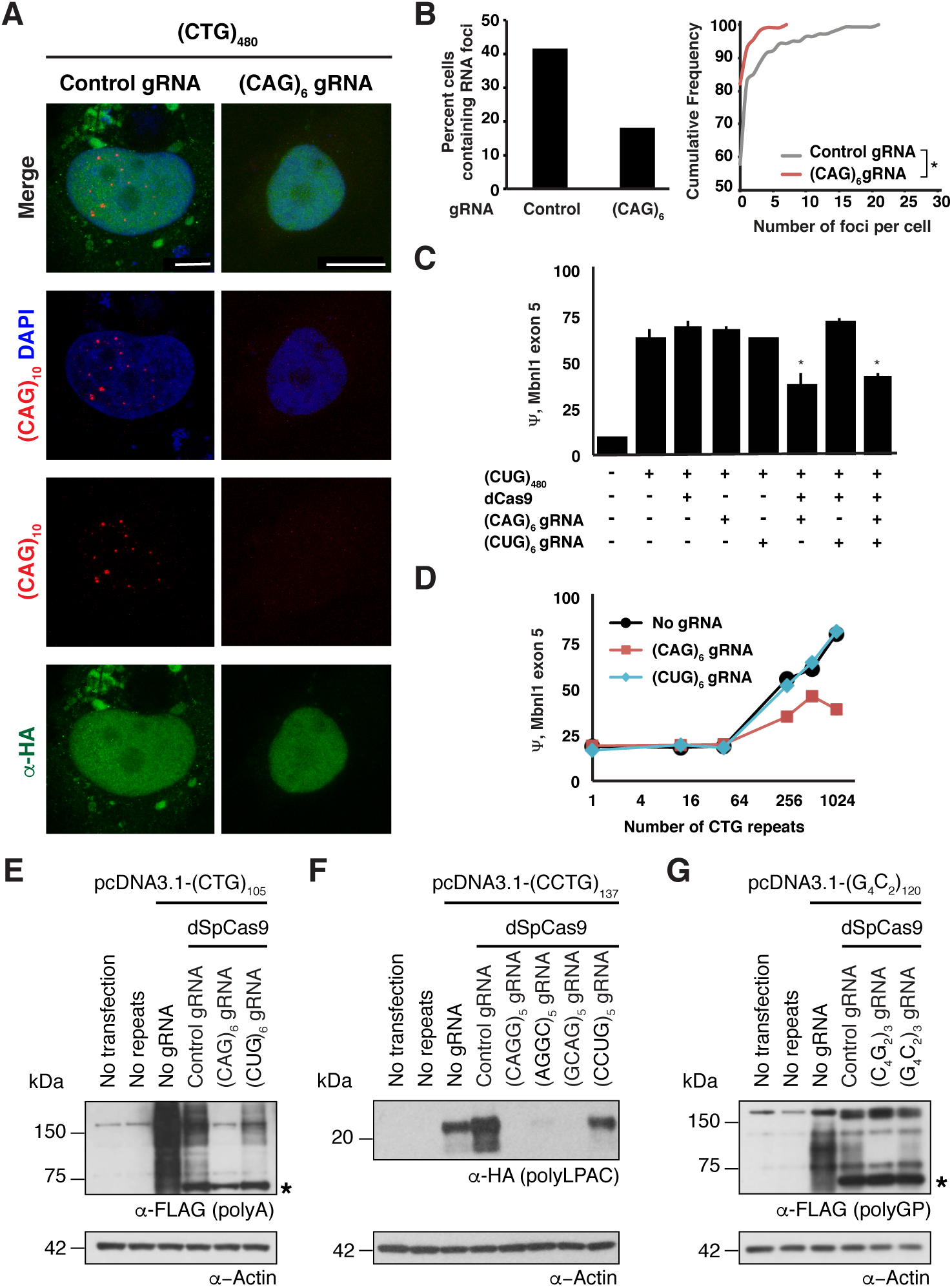
Transcriptional inhibition rescues molecular and cellular phenotypes in cell culture models of DM1, DM2, and C9ORF72/ALS/FTD. A) Representative images of HeLa cells transfected with plasmids expressing (CTG)_480_ repeats, dCas9 tagged with HA and control or (CAG)_6_ gRNAs. RNA foci are visualized by a (CAG)_10_ fluorescent oligonucleotide (red), dCas9 by an α-HA antibody (green), and DNA by DAPI (blue). Scale bar: 10μm B) Quantitation of the number of HA-positive cells showing nuclear RNA foci (left panel) and the cumulative density function of HA-positive cells with a given number of RNA foci (right panel), in the presence of dCas9 and control or (CAG)_6_ gRNAs (Kolmogorov-Smirnov test, *p < 0.005). C) Percent spliced in (Psi, Ψ) of MBNL1 exon 5 expressed in a minigene context in HeLa cells, in the presence of combinations of plasmids encoding (CTG)_480_ repeats, dCas9, and (CAG)_6_ or (CUG)_6_ gRNAs. N > 3 for all conditions (two-tailed T-test, *p < 0.005) D) Ψ of MBNL1 exon 5 expressed in a minigene context in HeLa cells, in the presence of plasmids encoding 0, 12, 40, 240, 480, or 960 CTG repeats, as well as dCas9 and (CAG)_6_ or (CUG)_6_ gRNAs. N > 3 for all conditions. Error bars are too small to be visible in this plot. E) Western blot against the FLAG-tagged poly-Ala RAN product expressed from DM1 CTG repeats transfected into HEK293T cells. Cells were transfected with various combinations of plasmids encoding no repeats or (CTG)_105_ repeats with combinations of dCas9 and control, (CAG)_6_ or (CUG)_6_ gRNAs. The RAN peptide migrates as a smear between 75 and 150 kD. The asterisk indicates protein produced by the dSpCas9 plasmid, which cross-reacts with the anti-FLAG antibody. F) Western blot against the HA-tagged LPAC RAN protein expressed from DM2 CCTG repeats transfected into HEK293T cells. Cells were transfected with various combinations of plasmids encoding no repeats or (CCTG)_137_ repeats with combinations of dCas9 and control, (CAGG)_5_, (AGGC)_5_, (GCAG)_5_ or (CCUG)_5_, gRNAs. The RAN peptide migrates around 20 kD. G) Western blot against the FLAG-tagged Poly-GlyPro RAN protein expressed from ALS G_4_C_2_ repeats transfected into HEK293T cells. Cells were transfected with various combinations of plasmids encoding no repeats or (G_4_C_2_)_120_ repeats with combinations of dCas9 and control, (C_4_G_2_)_3_ or (G_4_C_2_)_3_ gRNAs. The RAN peptide migrates as a smear between 75 and 120 kD. The asterisk indicates protein produced by the dSpCas9 plasmid, which cross-reacts with the anti-FLAG antibody. β-Actin serves as the loading control for A, B and C. N = 3 for A, B and C (representative blot shown).

The CTG, CCTG, and G_4_C_2_ repeats associated with DM1, DM2, and C9ALS/FTD, respectively, undergo repeat-associated non-ATG (RAN) translation^9^,^10^ (Zu et al, submitted 2017). We hypothesized that reduction in repeat-containing RNA abundance by dCas9-gRNA complexes would yield reduced RAN protein abundance. We assessed this by performing western analyses of RAN peptides robustly expressed from CTG, CCTG and G_4_C_2_ repeat containing plasmids transfected into HEK293T cells. Detection of the RAN products is facilitated by the presence of multiple tags downstream of the repeats, one in each coding frame (**Fig. 2E**). In all three western analyses, RAN proteins were observed only in the presence of repeat-containing plasmids, and were slightly reduced upon expression of a control, non-targeting gRNA, likely due to competition of plasmids for transcriptional machinery. Consistent with measurements of CUG-containing RNA levels (**Fig. 1B**), the FLAG-tagged poly-Ala peptide translated from CUG repeats was dramatically reduced in the presence of (CAG)_6_ gRNA, and modestly reduced in the presence of (CUG)_6_ gRNA (**Fig. 2E**). Importantly, the RAN protein was not reduced in the presence of (CAG)_6_ gRNA alone without dCas9 (**Fig. S2D**). HA-tagged poly-LPAC translated from CCUG repeats was dramatically reduced in the presence of (CAGG)_5_, (AGGC)_5_, and (GCAG)_5_ gRNAs (**Fig. 2F**), but not in the presence of (CCUG)_5_ gRNA (**Fig. S2E**). To test the effects of dCas9 on levels of poly-GlyPro translated from G_4_C_2_ repeats, we designed two gRNAs with NGG PAMs: (C_4_G_2_)_3_ that would target the non-template strand and (G_4_C_2_)_3_ that would target the template strand. Consistent with our studies of CTG and CCTG repeats, dCas9-gRNA complexes targeted to (G_4_C_2_)_120_ exhibited silencing of RAN peptide in a strand-dependent manner (**Fig. 2G**), and did not occur in the presence of gRNA alone (**Fig. S2F**). While reduction in RAN protein was achieved with both gRNAs, silencing was more effective with gRNA targeting the non-template strand.

### dCas9-mediated transcriptional inhibition reduces nuclear RNA foci and rescues splicing defects in human DM1 myoblasts

We next tested whether this approach could impede transcription of expanded repeats in the native *DMPK* locus, in primary human DM1 myoblasts. To express deactivated Cas9, we generated an adeno-associated virus (AAV) carrying sequences for the Cas9 and gRNAs that can efficiently infect muscle cells. As the traditional dCas9 sequence cannot be accommodated within the limits of the AAV genome, we turned to the smaller *S. aureus* Cas9 (SaCas9) that can be packaged with a U6 promoter-driven gRNA within the AAV genome^35^. We generated a deactivated SaCas9 (dSaCas9) by mutating catalytic residues D10 and H557 to alanines ^36^. Since dSaCas9 exhibits slightly different PAM preferences to dSpCas9 (NNGRR versus NGG), we used MBTA-Seq to confirm that dSaCas9 with a (CAG)_6_ gRNA could impede transcription of expanded CTG repeats in a length-dependent manner (**Fig. S3A**). Next, we packaged dSaCas9 with control or (CAG)_6_ gRNA into an AAV2/6 virus, as we found that the AAV6 capsid efficiently infects myoblasts in culture (**Fig. S3B**).

We infected a primary DM1 myoblast line with the AAV2/6 encoding dSaCas9 and (CAG)_6_ or control gRNAs. Expression of dSaCas9 protein with 5 nuclear localization signals in the infected cells was detected by immunofluorescence against the HA tag and was found to be nuclear, although at times also appeared to be cytoplasmic (**Fig. S3B**, **Fig. 3A**). We assayed CUG repeat content by FISH and as observed previously in HeLa cells, the number of CUG repeat foci per cell was decreased in the presence of dSaCas9 and (CAG)_6_ gRNA, relative to control gRNA (**Fig. 3A, B**).

**Figure 3.**
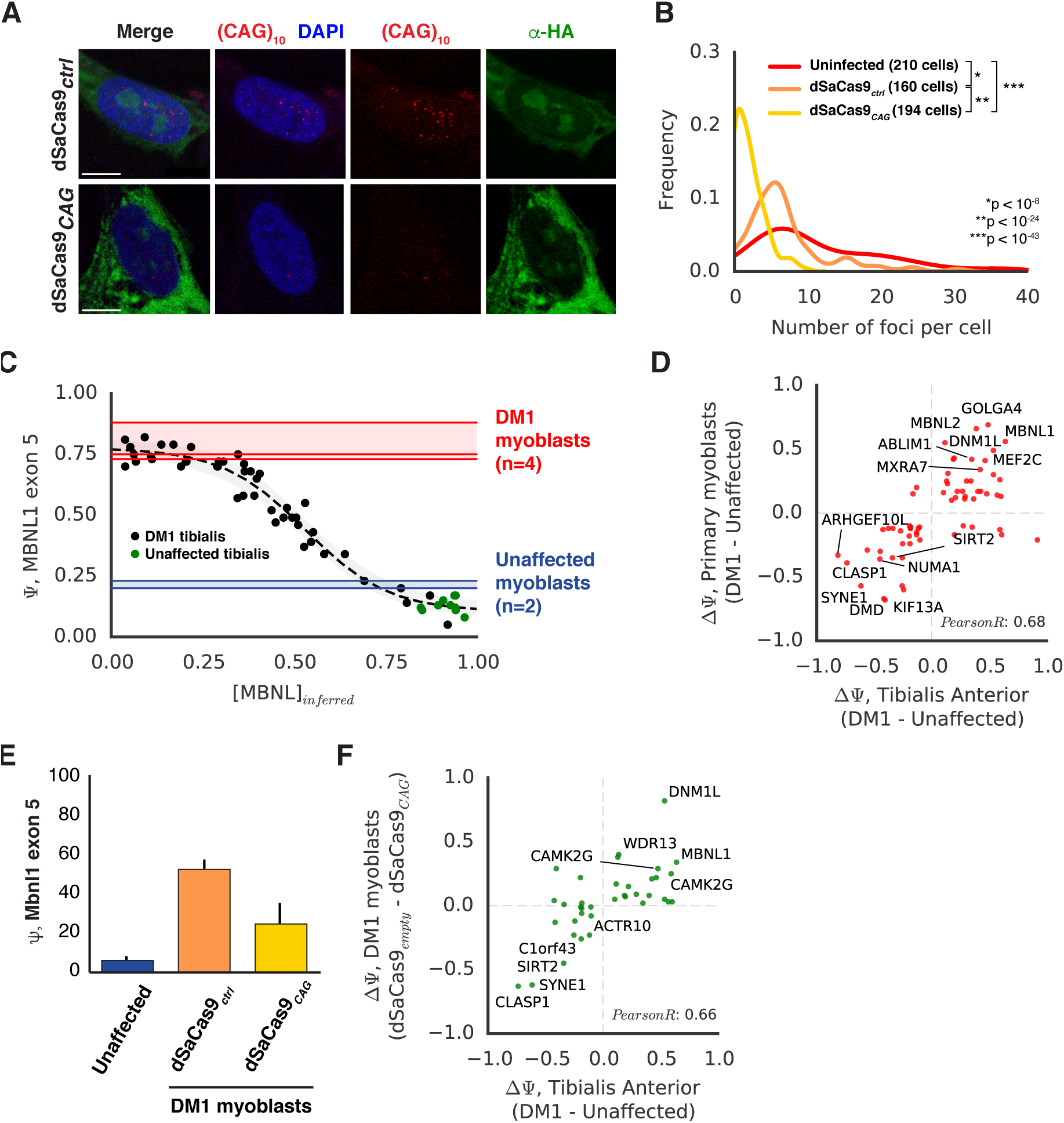
AAV-dSaCas9 rescues molecular and cellular phenotypes in human DM1 myoblasts. A) Representative images of DM1 myoblasts infected with AAV-dSaCas9 carrying control or (CAG)_6_ gRNAs. RNA foci are visualized by a (CAG)_10_ fluorescent oligonucleotide (red), HA-tagged dCas9 by an α-HA antibody (green), and DNA by DAPI (blue). Scale bar: 10μm B) Probability density function of cells with a given number of RNA foci, in the presence of control or (CAG)_6_ gRNAs. A Kolmogorov-Smirnov test was performed for statistical significance. C) MISO estimates of MBNL1 exon 5 Ψ in 11 unaffected and 44 DM1 tibialis anterior biopsies are plotted in order of [MBNL]_inferred_ as previously described. A sigmoid curve was fit to these points and shown (dashed line) with 90% confidence intervals (gray shading). MISO estimates of MBNL1 exon 5 Ψ values in an unaffected and DM1 primary myoblast line (2 and 4 replicates, respectively) are shown in blue and red lines, respectively, with ranges also shaded. D) Scatter plot for splicing events regulated in both tibialis biopsies and the human myoblast lines described in (C). 115 splicing events were selected from tibialis biopsies based on best sigmoid fits with [MBNL]_inferred_ as in (C). The x-axis of the scatter plot is the mean Ψ in the most severely affected biopsies (<0.33 [MBNL]_inferred_) minus the mean Ψ across unaffected individuals. The y-axis of the scatter plot is the mean Ψ in the DM1 myoblast minus the mean Ψ in the unaffected myoblast line. Labeled points have ΔΨ>0.3 in both conditions, a sigmoid fit<1.3, and a y-axis monotonicity score>1 (see Methods). Pearson correlation for all points is listed. E) MBNL1 exon 5 Ψ, assessed by RT-PCR in unaffected myoblasts and DM1 myoblasts infected with AAV-dSaCas9 carrying control or (CAG)_6_ gRNAs. F) Scatter plot illustrating changes in Ψ in response to AAV-dSaCas9 (CAG)_6_ gRNA, for splicing events exhibiting concordant behavior in (D). The x-axis is as in (D), but the y-axis is Ψ as assessed in the DM1 myoblast minus the mean Ψ in the unaffected myoblast line. Labeled points have ΔΨ>0.1 in both conditions and a y-axis Bayes Factor>5 (see Methods). Pearson correlation for all points is listed.

Although splicing events in DM1 patient muscle have been well characterized, myoblasts in culture do not express many transcripts typically present in mature muscle. To identify molecular changes in cultures that appropriately model *in vivo* DM1 biology, we further characterized the line described above, as well as a line derived from an individual unaffected by DM1. Four independent RNAseq libraries were created from the DM1 cells grown in several different growth conditions, and two from the unaffected line. To identify splicing changes in these lines that appropriately model those occurring in human DM1 muscle, we first defined a high confidence set of DM1-relevant splicing events by re-analyzing a set of 55 transcriptomes (44 DM1 and 11 unaffected) from human tibialis biopsies (GSE86356). We identified splicing events whose inclusion level, Ψ, strongly correlated to the concentration of free, functional MBNL protein in the affected tissue, as previously described^34^. We fit sigmoid curves describing the relationship between MBNL concentration and Ψ, for example for MBNL1 exon 5 (**Fig. 3C**). The best-fitting events were selected (see Methods, **Fig. S4**), and their dysregulation (ΔΨ, Tibialis anterior, DM1 minus unaffected) was plotted against dysregulation observed in the DM1 myoblasts (ΔΨ, Primary myoblast, DM1 minus unaffected) (**Fig. 3D**). These filtering steps yielded a set of events whose behavior is reasonably recapitulated in each system; the correlation in splicing dysregulation between tibialis and myoblasts was ~0.68.

To assess splicing rescue in DM1 myoblasts upon treatment with AAV-dSaCas9 and (CAG)_6_ gRNA, we first performed RT-PCR to measure the inclusion level of MBNL1 exon 5. While viral infection with an AAV encoding dSaCas9 and control gRNA yielded a Ψ of ~50%, infection with AAV encoding dSaCas9 and (CAG)_6_ gRNA rescued Ψ to ~20% (**Fig. 3E**). We assessed splicing transcriptome-wide by RNAseq, and analyzed those exons in the myoblasts whose changes are concordant with changes observed in tibialis biopsies (lower left and upper right quadrants, **Fig. 3D**), and whose baseline Ψ values are less than 0.33 units apart when comparing tibialis to myoblasts. Several exons were successfully rescued, and a correlation of ~0.66 was observed between dysregulation in tibialis (ΔΨ, DM1 minus unaffected) versus rescue by dSaCas9 and (CAG)_6_ gRNA (ΔΨ, control gRNA minus (CAG)_6_ gRNA) (**Fig. 3F**, **Table S3**). To assess potential off-target changes in gene expression for other transcripts with genomic CTG or CAG repeat tracts, we treated non-DM1 myoblasts with AAV-dSaCas9 and (CAG)_6_ gRNA or control gRNA and performed RNA-Seq. We enumerated the longest contiguous CTG or CAG repeat tracts in all pre-mRNAs in the human genome, and found none to exceed 24 repeats; maximums were 24 CTG in TCF4 and 22 in AR. We observed no differences in gene expression that depended on the length of repeat tracts, for both CTG or CAG repeats (**Fig. S3C, D**).

These observations suggest that dSaCas9 targeted to CTG repeat tracts can impede transcription of expanded CUG repeats, relieve MBNL sequestration, and restore splicing homeostasis in human cells containing DM1 repeat expansions, with selectivity that depends on the extreme repeat lengths commonly found in symptomatic tissue.

### dCas9-mediated transcriptional inhibition reduces RNA foci, rescues Clcn1 expression, and decreases myotonia in a mouse model of DM1

We next assessed whether dCas9 could impede transcription of expanded CTG repeats in a well-established mouse model of DM1, *HSA*^*LR*^ ^37^. These mice carry a human skeletal actin transgene containing 250 CTG repeats in the 3’ UTR and exhibit molecular, cellular, and phenotypic properties characteristic of DM patients. To assess the efficacy of this approach independently of potential *in vivo* drug delivery issues, we first dissected extensor digitorum longus muscle (EDL) fibers, and cultured them *ex vivo.* As CTG repeat expression in this model is driven by the *HSA* promoter, RNA foci are ubiquitous and numerous in myonuclei (**Fig 4A**, **Top panel**). To assess the extent to which dSaCas9 could reduce RNA foci, we infected cultured individual *HSA* EDL fibers for 2 days and performed RNA FISH against CUG repeats (**Fig. 4A**). In untreated *HSA* fibers, we observed robust FISH signal in ~80% of myonuclei. This remained at similar levels in the presence of a dSaCas9 with control gRNA, but was reduced to ~50% in the presence of dSaCas9 with (CAG)_6_ gRNA, where any sign of FISH signal in a nucleus was recorded (**Fig. 4B**). Overall FISH signal intensity, quantitated in >1400 nuclei per condition (see Methods, **Fig. S5**), was dramatically decreased in fibers treated with dSaCas9-(CAG)_6_ gRNA relative to control gRNA and untreated fibers (**Fig. 4C**). These observations suggest that efficient delivery of dSaCas9-(CAG)_6_ to muscle fibers *ex vivo* is sufficient to significantly reduce RNA foci load within 2 days.

**Figure 4.**
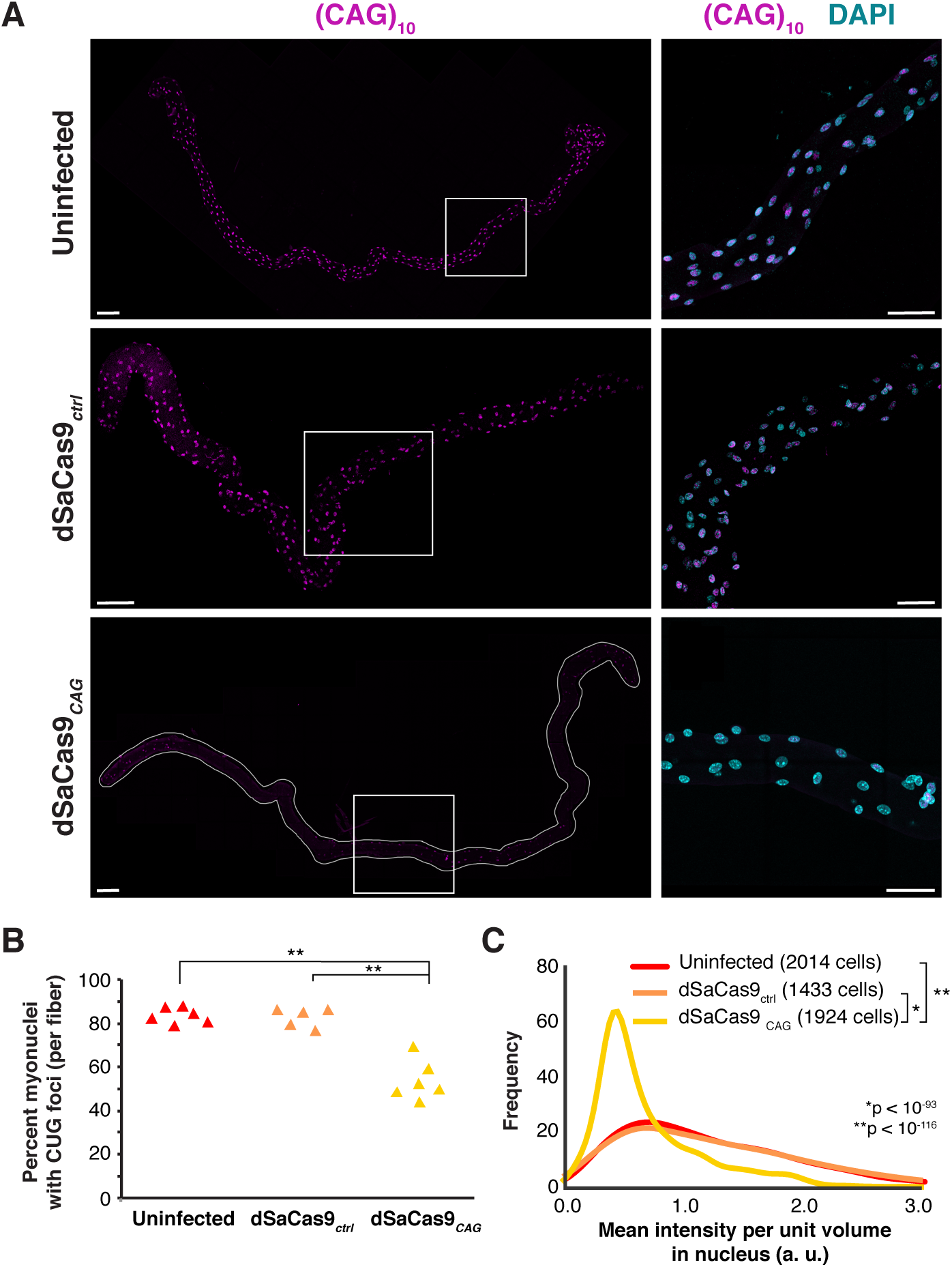
AAV-dSaCas9 reduces RNA foci in *HSA*^LR^ mouse muscle fibers. A) FISH to detect nuclear RNA foci by a fluorescent oligonucleotide (CAG)_10_ probe (magenta) in the myonuclei of EDL muscle fibers from the *HSA*^LR^ mouse that were untreated (top), infected with AAV-dSaCas9-control gRNA (middle) or AAV-dSaCas9-(CAG)_6_ gRNA (bottom). One representative fiber for each condition is shown on the left (Scale bar: 100μm) and insets from each fiber (white boxes) are shown on the right (Scale bar: 50μm). DAPI is in cyan. B) The percentage of myonuclei per fiber showing RNA foci was quantitated across all 3 conditions with 5-6 fibers per condition. Each fiber contains 400-500 nuclei in all cases (two-tailed T-test, **p < 0.0005). C) Probability density function of intensity of FISH signal in myonuclei from untreated fibers (red), fibers infected with AAV-dSaCas9-control gRNA (orange) or AAV-dSaCas9-(CAG)_6_ gRNA (yellow). A Kolmogorov-Smirnov test was performed for statistical significance.

Having observed robust elimination of RNA foci *ex vivo*, we turned to *in vivo* experiments. Given previous reports of immune reactivity against Cas9, especially following intramuscular injection we chose to administer AAV-dSaCas9-gRNA by temporal vein injection at postnatal day 2, prior to full establishment of immune tolerance. AAV2/6 or AAV2/9 carrying dSaCas9 and (CAG)_6_ or control gRNA was injected. Five weeks following injection, electromyography was performed to analyze myotonia in tibialis anterior and gastrocnemius muscles (**Fig. 5A**). Wild type FVB mice showed no myotonia, an *Mbnl1* KO mouse showed myotonia in 100% of needle insertions, and *HSA*^LR^ mice showed myotonia in 87-100% of insertions. *HSA*^LR^ injected with AAV6-dSaCas9-control gRNA showed myotonia levels similar to uninjected *HSA*^LR^ (87%). However, *HSA*^LR^ injected with AAV6-dSaCas9-(CAG)_6_ gRNA showed a reduction in myotonia, with some mice showing myotonia in 33-50% of insertions. Interestingly, nearly all mice injected with AAV2/6 showed rescue while only 1 mouse injected with AAV9 showed rescue (**Fig. S5D**).

**Figure 5.**
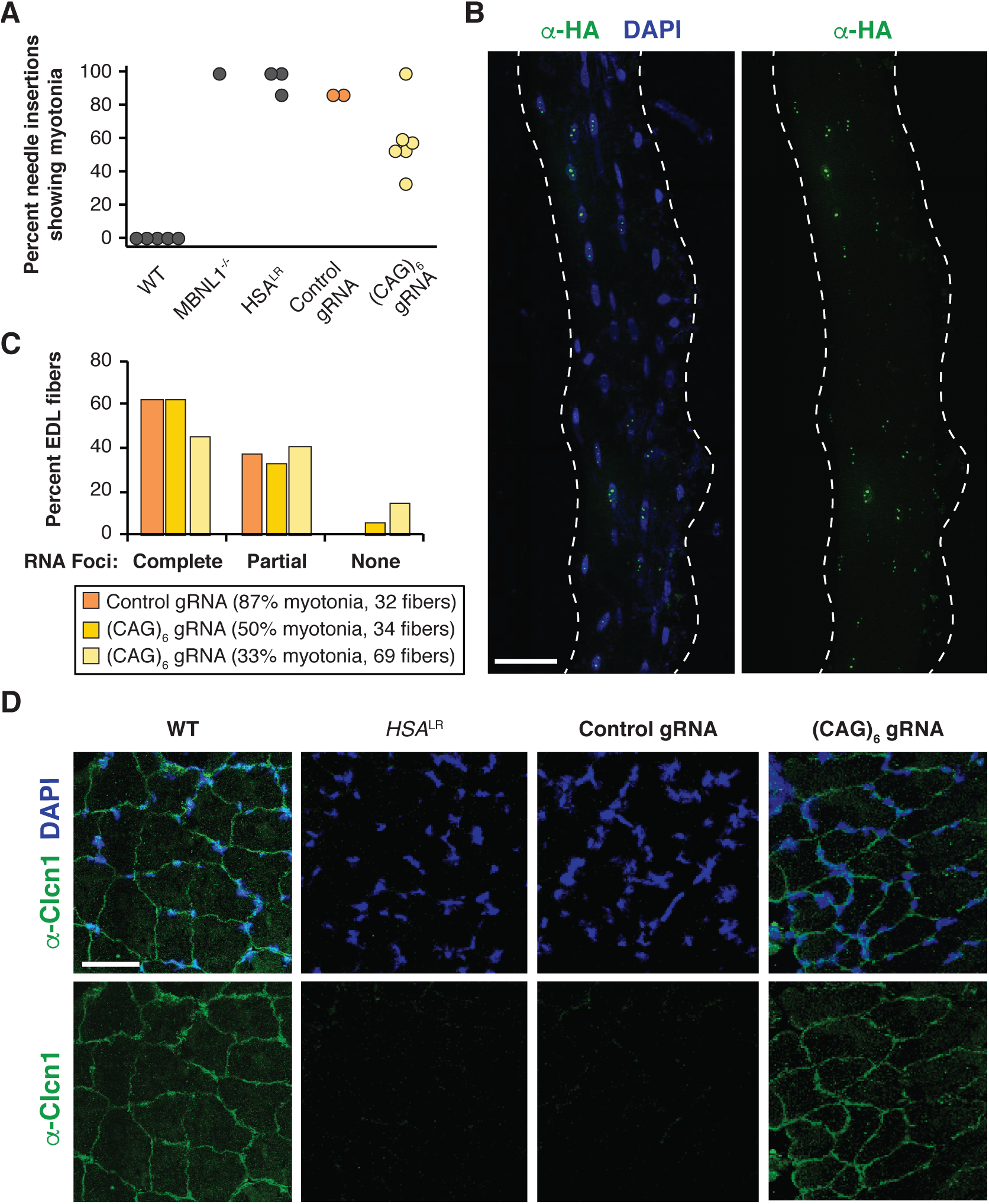
AAV6-dSaCas9-(CAG)_6_ rescues muscle phenotypes in *HSA*^LR^ mice. A) Percent needle insertions showing a myotonic run in wild type FVB, MBNL1^-/-^, *HSA*^LR^, (black circles) and *HSA*^LR^ treated with AAV6-dSaCas9 with control gRNA (orange circles) or (CAG)_6_ gRNA (yellow circles). Each point represents a muscle from a single animal, but in some animals, both tibialis anterior and gastrocnemius were assayed. N=2 mice for control gRNA and N=4 mice for (CAG)_6_ gRNA. B) Immunofluorescence against HA-dSaCas9 in a single EDL fiber from a mouse treated with AAV2/6-dSaCas9 and (CAG)_6_ gRNA. DAPI is also shown in the merged image. Scale bar, 50 μM. C) Quantitation of numbers of EDL fibers showing complete, partial, or no CUG RNA foci upon treatment with AAV2/6-dSaCas9 and control gRNA or (CAG)_6_ gRNA. Myotonia levels and numbers of fibers analyzed are also listed. D) Immunofluorescence against Clcn1 in tibialis muscle sections from WT FVB, *HSA*^LR^, and *HSA*^LR^ treated with AAV-dSaCas9 and control gRNA or (CAG)_6_ gRNA. DAPI is shown in blue. Scale bar, 50 μM.

Because mis-splicing of chloride channel 1 (Clcn1) is well-established to mediate myotonia in DM1 and in *HSA*^LR^ mice, we measured splicing patterns of Clcn1 mRNA in treated mice. Surprisingly, when analyzing RNA extracted from bulk muscle tissue, we could not detect significant rescue in Clcn1 splicing, or other MBNL-dependent splicing events (data not shown). Because myotonia is measured by assaying individual bundles of fibers, we hypothesized that some regions of muscle may be rescued and not others, and that any potential change in isoform composition in these regions may be diluted in analyses of bulk tissue. This is consistent with a roughly ~2.5-fold increase in myonuclei per fiber over the first 4 weeks of post-natal development, which could also dilute the proportion of myonuclei containing AAV episomes^39^. AAV6-delivered dSaCas9 distribution in EDL fibers, as assessed by immunofluorescence against the HA tag, revealed mosaic expression, with rare fibers showing region-specific nuclear signal for dCas9 (**Fig. 5B**), and most fibers not showing presence of dCas9 at all. Control immunofluorescence experiments against lamin A showed ubiquitous and consistent labeling of all myonuclei, ruling out potential staining artifacts (**Fig. S5E**). In spite of mosaic dSaCas9 distribution, 5-15% of fibers showed complete loss of CUG RNA foci (**Fig. 5C**). Furthermore, animals that showed the greatest decrease in RNA foci also showed the strongest myotonia rescue. Reduction of RNA foci load in some nuclei raised the possibility that proper Clcn1 splicing could be achieved in a subset of nuclei, and that these mRNAs may spread locally throughout the fiber to produce functional Clcn1 protein. Indeed, mice treated with (CAG)_6_ gRNA showed increased Clcn1 staining at muscle membranes relative to mice treated with control gRNA, but Clcn1 protein was localized only to a subset of fibers per muscle section. These observations are consistent with observations of myotonia elimination in a subset of fibers, yet absence of splicing rescue when assessing bulk tissue (**Fig. 5D**). In summary, these experiments suggest that dCas9 can, in principle, rescue disease phenotypes via transcriptional repression, but that widespread rescue of molecular events in muscle will require efficient delivery to a large proportion of myonuclei expressing toxic RNA.

## Discussion

The application of CRISPR/Cas9 to genetic diseases is thought to hold great therapeutic promise. Multiple groups have successfully edited genomes in cell and animal models of disease. Some have directly applied these approaches to microsatellite expansion diseases to remove the expanded repeat tract^40^ or cause somatic instability leading to repeat contraction^41^. However, challenges remain to ensure both efficient and specific cleavage activity across multiple cells and tissues of the body. Here, we explore an alternative form of Cas9, dCas9, in which cleavage activity is abrogated. Previous studies demonstrated that dCas9 can successfully reduce transcript levels in prokaryotes and this effect is achieved primarily by blocking elongation of RNA Pol II through the gene body rather than targeting the transcripts themselves^22^. In eukaryotes, however, transcriptional inhibition by recruiting dCas9/gRNA complexes to the gene body was inefficient. As expanded microsatellites are thought to be more challenging for RNA Pol II elongation, we hypothesized that dCas9/gRNA recruitment would effectively impede elongating polymerase. By testing disease-associated repeat sequences across multiple repeat lengths *in vitro* and in disease models, we demonstrate that dCas9 can substantially reduce repeat-containing transcript abundance. Importantly, repression efficiency is proportional to repeat length, because longer repeats recruit an increased number of dCas9/gRNA complexes, leading to greater transcriptional blockade (**Fig. 1B-C**).

dCas9 is well established to target DNA^22^, but has also been shown to bind repeat-containing RNA in a gRNA-dependent manner^42^. Indeed, when we immunoprecipitated dCas9 and assessed RNA binding, we observed increased binding to transcripts containing longer repeats (**Fig. S2G**). However, as repeat length increases, the amount of successfully transcribed RNA decreases, suggesting that at long repeat lengths, less RNA remains to be targeted. In the context of disease, microsatellite repeat tracts undergo dramatic somatic expansion in DM1, DM2, C9ALS/FTD, and HD, and evidence suggests that repeat lengths in tissues of symptomatic individuals reach thousands of nucleotides in length^2^,^3^. Therefore, DNA-based targeting by dCas9 may play a primary role in driving potential therapeutic benefits in individuals with full expansions (**Fig. 6**). At short and intermediate repeat lengths, therapeutic benefit may derive from both DNA-and RNA-based targeting by dCas9, to not only impede transcription of repeat-containing DNA, but also modulate downstream consequences of transcribed “escaper” RNAs. Taken together, we demonstrate that dCas9 is an effective tool to limit downstream effects of toxic RNAs and present proof of concept data suggesting that therapeutic benefit can be achieved through this approach.

**Figure 6.**
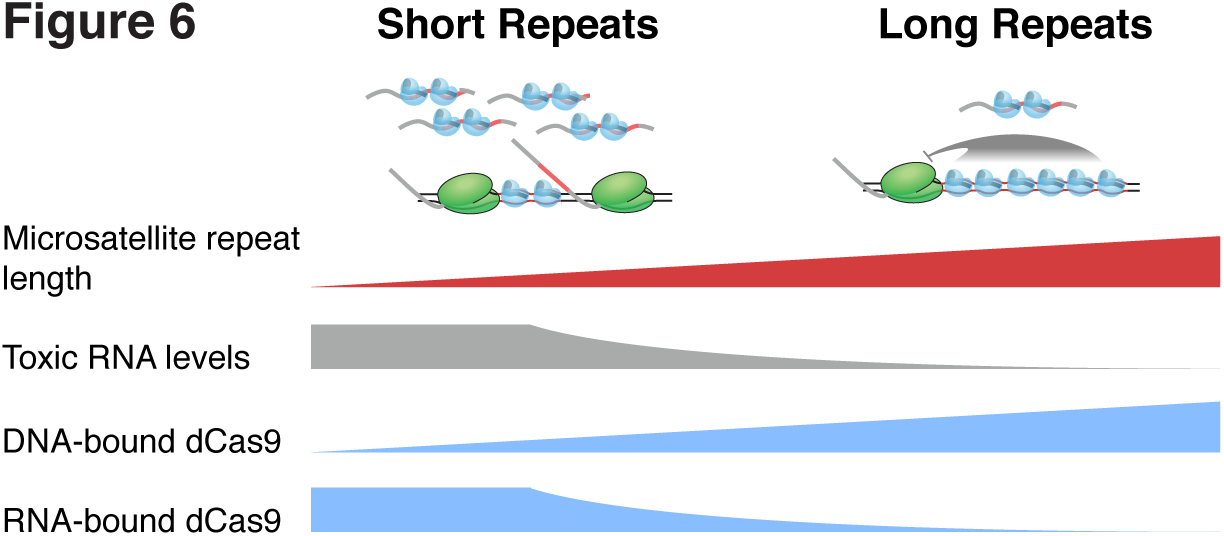
Model for microsatellite repeat expansion targeting by dCas9. dCas9 binding to DNA impedes transcription of long, expanded microsatellite repeats. In the context of shorter repeats, fewer copies of DNA-bound dCas9 may be insufficient to fully inhibit elongation by RNA polymerase II, permitting production of RNA that can also be targeted by dCas9.

MBTA-Seq allowed us to rapidly and accurately measure expression of transcripts that are traditionally difficult to assay, in particular across repeat types and repeat lengths in a highly controlled manner. In principle, this approach can be adapted for any collection of transcripts, in the context of transient transfections or stable cell lines. We used this assay to quantitatively measure how PAM sequence and target strand influence the efficacy of transcriptional blockade by dCas9; these effects likely directly relate to the strength of binding by Cas9 as well as the ability of RNA Pol II to unwind DNA. Although we show here that CTG, CCTG, and G_4_C_2_ repeats are targeted effectively by dCas9, in principle, alternative Cas9 variants favoring PAM sequences encoded on the non-template strand of microsatellite repeats should also be effective. Importantly, by targeting gRNAs with similarly effective PAMs to template versus non-template strand transcripts, dCas9 can be used to separate effects arising from sense versus antisense repeat-containing transcripts, such as those occurring in C9ALS/FTD.

Effective, safe delivery in a multi-systemic fashion to sufficient numbers of post-mitotic cells will be necessary for this approach to be therapeutically viable in the context of many of these dominantly inherited neurological diseases. In DM and C9ALS/FTD, it is unclear what fraction of nuclei must be rescued in multi-nucleated muscle or in the central nervous system to halt or improve disease symptoms. Here, we observed that rescue of a subset of nuclei was sufficient to yield partial rescue of myotonia. However, myotonia can be modeled as a loss-of-function event in which Clcn1 protein is lost, and there is evidence that in other recessive muscle diseases, gene editing of a small subset of nuclei can restore sufficient protein expression across muscle fibers to yield therapeutic benefit^43^,^44^. Conversely, mis-splicing events yielding pathogenic isoforms with dominant-negative behavior may require full elimination to mitigate deleterious consequences.

Our approach used systemically delivered AAV, which revealed regional variation in transduction efficiency, likely because NLS-tagged Cas9 protein remained restricted to myonuclear domains transduced by AAV episomes, and many new myonuclei are recruited to fibers throughout the first 4 weeks of post-natal muscle development^39^. It is possible that more localized or concentrated delivery would have facilitated transduction of a higher proportion of myonuclei. Previous studies with AAV-mediated gene therapy typically evaluate expression of proteins that can spread throughout muscle fibers, precluding measurement of transduction efficiency to all myonuclei. Regional variation in rescue is likely to be less relevant with therapies with more homogeneous delivery, for example, small molecules. Nevertheless, reduction of toxic RNA repeat load in a subset of nuclei was sufficient to increase production of Clcn1 protein, with consequences for distant regions of muscle naïve to dCas9. These results emphasize how differences in mechanism of action and delivery efficiency between various therapeutic approaches should significantly influence how we interpret changes in response to those therapies, both at the molecular and phenotypic level.

Finally, while this approach mitigates a key safety concern of CRISPR, which is the unwanted cleavage of off-targets, it also raises questions about whether long-term exposure to dCas9 can be achieved and tolerated, a necessity for repeat expansion diseases. Perhaps most importantly, the ability of dCas9 to impede transcription of microsatellite expansions defines a window in which transcriptional inhibition of expanded repeats is feasible, yet does not interfere with that of typical RNAs. This approach may serve as a baseline with which to compare alternative therapeutic approaches, as well as a tool to identify mechanisms and principles by which transcriptional activity of RNA polymerase II can be modulated in a sequence-specific manner.

## Materials and Methods

### Cell culture, cell lines and transfection

HeLa and HEK293T cells were cultured in 1X DMEM supplemented with 10% fetal bovine serum and 1% Penicillin/Streptomycin at 37°C and 5% CO_2_. The DM1 primary myoblast cells were obtained by G. Xia. The clinical data of research subjects from who muscle biopsies were obtained, have been described before^45^. Muscle biopsies were performed using 7G UCH Muscle Biopsy Needle (#8066, www.cadencescience.com). Samples were cut into small pieces and seeded into a 6cm dish. Myoblasts were expanded at a 1:2 ratio in myoblast growth media (Skeletal Muscle Cell Growth Medium-2, Lonza, #CC 3245) at 37°C and 5% CO_2_. For viral transductions, these primary myoblasts were maintained in the growth medium for 3 days and then switched to Differentiation media (DMEM-F12 supplemented with 2% horse serum) for 3 days.

Plasmid DNA transfections of HeLa and HEK293T cells were performed using Trans-IT LT1 (MirusBio) as per manufacturer’s instructions. For RNA and FISH analyses, HeLa cells were harvested or processed 72 hours post transfection.

### Cloning of barcoded repeat plasmids and gRNA constructs and generation of stable repeat expressing HeLa cell lines

Non-barcoded plasmids carrying 0, 12, 40, 240, 480, 960 CTG repeats (CTG_n_) and 960 CAG repeats (CAG_960_) were kind gifts from Tom Cooper (Baylor College of Medicine). Plasmids with 0 and 12 repeats were modified so that their vector backbones were identical to the others, and carried the ampicillin resistance gene. The (CCTG)_240_ plasmid was created as in Philips et al ^46^. In brief, oligonucleotide fragments 5′-TCGA(CCTG)_20_C-3′ and 5′-TCGAG(CAG)_20_-3′, were phosphorylated, annealed, gel isolated, and concatemerized by T4 DNA ligase. Concatemers not in a head to tail orientation were digested by SalI and XhoI. Concatemers were gel-isolated and cloned into the SalI site of CTG_0_. To create CAGG_240_, CTG_960_ was digested with SalI and HindIII and a new fragment was introduced which included HindIII and AgeI restriction sites (DT_MCS). CCTG_240_ was digested with XmaI and HindIII and ligated with DT_MCS digested with HindIII and AgeI to reverse orientation of the repeats and form CAGG_240_.

For MBTA-Seq, each repeat-containing plasmid was barcoded by introducing a fragment containing a random 8-nt sequence at the PflMI restriction site located downstream of the repeats via In-Fusion cloning (Clontech). Clones were sequenced to confirm that each repeat containing plasmid carried a unique barcode (Fig S1A). Barcoded CTG_960_ and CAG_960_ plasmids were generated by digesting (CTG)_960_ and (CAG)_960_ with HindIII and AccIII and ligating gBlocks containing unique barcodes with the digested plasmids.

To generate stable cell lines, the *DMPK* expression cassette was removed from each of the 6 barcoded plasmids and inserted into pAC156 (gift from Albert Cheng), a plasmid with Piggybac transposon terminal repeats as well as a puromycin selection cassette. All 6 plasmids were transiently transfected together with the Piggybac mPB transposase into HeLa, and selected by puromycin. Single cells were isolated by flow cytometry, and colonies were cultured in 96 well plates. 48 colonies were expanded and subjected to MBTA-Seq to screen for integration and expression of all 6 plasmids. Counts for all barcode deep sequencing experiments can be found in Table S1.

The dCas9 gRNAs were cloned into an AflII-digested U6 expression vector by annealing oligos as previously described^47^. The dSaCas9 gRNAs were cloned into BsaI - digested vector by annealing oligos as previously described^35^.

### dCas9 chromatin and RNA IP

dCas9 ChIP and RIP experiments were performed on the HeLa cell line containing the six different repeat lengths (CTG0-960) integrated into genome. 10cm plates of ~80% confluent cells were transfected with the pXdCas9 and U6 expression vectors expressing the dCas9 protein and the control or (CAG)_6_ gRNAs respectively ^4847^, using the manufacturers guidelines. For ChIP, 48 hours after transfection cells were crosslinked with 1% formaldehyde for 20 minutes at room temperature. Cross-linking reactions were stopped by addition of glycine to a final concentration of 0.125 M. Cells were then harvested, washed with phosphate buffered saline and pelleted. 1 mL of Lysis Buffer 1 (50mM Hepes [pH 7.5], 140mM NaCl, 1mM EDTA, 10% glycerol, 0.5% Igepal, 0.25% Triton X-100) was added to the cells and rocked at 4°C for 10 minutes. After spinning, the cells were incubated in Lysis Buffer 2 (10mM Tris-HCl [pH 8.0], 200mM NaCl, 1mM EDTA, 0.5mM EGTA) for 10 minutes at RT. Nuclei were pelleted and resuspended in 1mL of Lysis Buffer 3 (10mM Tris-HCl [pH 8.0], 100mM NaCl, 1mM EDTA, 0.5mM EGTA, 0.1% Na-deoxycholate and 0.5% N-lauroylsarcosine) and subjected to sonication in a Covaris S220 to obtain DNA fragments averaging 4kb in length. One-twentieth of the total chromatin served as input. The remaining material was used in the IP, which was performed using the HA-tag (C29F4) rabbit mAb conjugated to magnetic beads (Cell Signaling Technologies) at 4°C overnight to pull down the HA-tagged dSpCas9 and interacting DNA. Beads were washed 7 times with wash buffer (50mM Hepes [pH 7.6], 500mM LiCl, 1mM EDTA, 1% Igepal and 0.7% Na-Deoxycholate). Immunocomplexes were eluted from the beads with elution buffer (50mM Tris-HCl, 10mM EDTA and 1% SDS) at 65°C for 15 minutes. Crosslinks in the IP and input were reversed overnight at 65°C and treated with RNase A and proteinase K to remove RNAs and proteins. DNA was extracted with phenol-chloroform and precipitated with ethanol. The barcoded region associated with each repeat length was amplified from the isolated DNA fragments using primers containing adapters facilitating deep-sequencing.

For the RNA IP experiments, 48 hours after transfection, cells were harvested in 1mL of lysis buffer (100mM Kcl, 5 mM MgCl2, 10mM HEPES [pH 7.0], 0.5% Igepal, 1mM DTT, 100 U/mL SUPERase In RNase inhibitor (Thermo Fisher), 2mM vanadyl ribonucleoside complexes solution, 25uL/mL protease inhibitor cocktail). One-twentieth of the resulting lysate was used as input and the remaining lysate was incubated with HA-tag (C29F4) rabbit mAb conjugated to magnetic beads (Cell Signaling Technologies) at 4°C overnight to pull down the HA-tagged dSpCas9 and interacting RNA. Beads were washed with the lysis buffer four times at 4°C and immunocomplexes were eluted off with 0.1% SDS and proteins removed using proteinase K at 50°C for 30 minutes. RNA was isolated using the Direct-zol RNA miniprep kit (Zymo Research) and contaminating DNA was eliminated using TURBO DNase (Thermo Fisher). cDNA was generated using Superscript IV Reverse Transcriptase (Thermo Fisher) and subsequently barcoded regions were amplified using flanking primers carrying sequences suitable for deep sequencing.

### FISH and immunofluorescence analyses

To detect nuclear RNA foci, cells or muscle fibers were fixed with 4% PFA for 10 minutes at room temperature followed by ice cold RNAse free 70% ethanol for 30 mins. Fixed samples were washed with a 25% formamide wash buffer at 30°C for 30 mins and then hybridized with a CalFluor 610 conjugated (CAG)_10_ oligonucleotide (Biosearch Technologies) in a 25% formamide hybridization buffer overnight at 30^°C^. Finally samples were washed two times at 30°C with wash buffer for 30 minutes to an hour, incubated with DAPI (1mg/mL) and mounted in Vectashield.

Further IF analysis was performed on HeLa cells and DM1 myoblasts to detect the presence of the dCas9-HA protein. After excess oligonucleotide was washed off, the cells were blocked in 3% normal goat serum in 1% Triton X-100-PBS for 30 mins at room temperature, incubated with anti-HA antibody (1:500, #3724, Cell Signaling Technologies) overnight at 4^°C^, washed with 1X PBS, incubated with Alexa Fluor 488 conjugated anti-rabbit secondary antibody (1:500, Life Technologies) for 2 hours, washed and incubated with DAPI for 5 mins and mounted in Vectashield. Slides were imaged using the Zeiss LSM 880 Confocal Laser Scanning Microscope. IF analysis to detect dCas9 in mouse muscle fibers from the EDL of mice injected with AAV6-dSaCas9-(CAG)_6_ was conducted as described above, except samples were fixed with 100% isopropanol at −20°C for 10 mins.

Clcn1 was detected in mouse muscle by performing IF on frozen muscle sections of the TA. Frozen muscle was sectioned into 10uM slices, fixed with 100% acetone at −20°C for 20 minutes, washed with 0.3% Triton X-100-PBS and incubated with rabbit anti Clcn1 (1:100, #CLC11-S, Alpha Diagnostic International) overnight at 4^°C^. Samples were incubated with goat anti-rabbit Alexa Fluor 568 (1:500, Thermo Fisher) for 2 hours at RT and then treated with DAPI and mounted in Vectashield.

### Western blots for RAN peptides

HEK293Tcells were transfected with one of the following plasmids that express tagged RAN translated products: pcDNA-6XStop-(CTG)_150_-3X(FLAG-HA-cMyc-His), pcDNA-6XStop-(CCTG)_137_-3X(FLAG-HA-cMyc-His) or pcDNA-6XStop-(G_4_C_2_)_120_-3X(FLAG-HA-cMyc-His)^9^,^10^ (Zu et al, submitted 2017). These cells were co-transfected with the pXdCas9 plasmid expressing the dCas9 protein^48^, and U6 expression vectors^47^ expressing the control gRNA or gRNA’s targeting either strand of the CTG, CCTG and G_4_C_2_ repeats ((CAG)_6_ or (CUG)_6_ gRNAs, (CAGG)_5_, (AGGC)_5_, (GCAG)_5_,or (CCUG)_5_ gRNAs and (C_4_G_2_)_3_ or (G_4_C_2_)_3_ gRNAs, respectively).

72 hours after transfection, cells in each well of a 12-well tissue culture plate were gently rinsed 1X with PBS and lysed in 200ul of RIPA Buffer (50 mM Tris-Cl pH 7.4, 150 mM NaCl, 0.1% Na-Deoxycholate, 1% NP-40, 0.5% SDS) with protease inhibitors for 30 minutes on ice. Genomic DNA was sheared by 8-10 passages through a 21-gauge needle. The resulting lysate was centrifuged at 18,000 x g for 15 min and the supernatant was collected. The protein concentration of the lysate was determined using Pierce™ BCA Protein Assay. Equal amounts of protein were loaded and separated on a 4-12% NuPage Bis-Tris gel (Novex) and transferred to a nitrocellulose membrane (Amersham). The membrane was blocked in 5% milk in PBS-Tween20 (0.05%) for 1 hour and probed with anti-FLAG (1:2000) or anti-HA (1:1000) antibody in 1% milk solution in PBS-Tween20 (0.05%) overnight at 4°C. After the membrane was incubated with anti-mouse and anti-rabbit HRP (1:10,000) for 2 hours at room temperature, the bands were detected using the SuperSignal™ West Femto Maximum Sensitivity Substrate as per manufacturers protocol ^9^.

### Recombinant AAV production

Viral production was achieved through transfection of HEK293T cells cultured in 150mm plates with the pAAV6 serotype packaging plasmid^49^, pXX6 helper plasmid that contains the adenovirus E4, VA and E2a helper regions^50^ and AAV2-ITR containing plasmid expressing dSaCas9 and the control or (CAG)_6_ gRNA (generated from the SaCas9 plasmid pX601-AAV-CMV∷NLS-SaCas9-NLS-3xHA-bGHpA;U6∷BsaI-sgRNA, Addgene #61591). Transfections were carried out using the TransIT-LT1 transfection reagent and recommended protocols (MirusBio). Cells were harvested between 48h and 72h post-transfection, recombinant AAV2/6 virus was purified by iodixanol step gradients followed by vector concentration and buffer exchange with lactated Ringer’s in an Apollo 150kDa concentrator (Orbital Biosciences)^51^. Virus titers were determined using the Quant-iT Picogreen dsDNA assay kit (Life Technologies)^52^ and found to be ~10^11^ vg/mL.

### Isolation of single muscle fibers

Single muscle fibers were isolated from 3-4 week old *HSA^LR^* mice as described previously^53^. Briefly, the EDL was dissected and digested with a 0.2% Collagenase Type I in DMEM solution in a 37°C water bath for 1 hour without agitation. The digested muscle was then flushed with DMEM to separate out individual muscle fibers. Fibers were cultured in DMEM containing 20% FBS overnight before infection with AAV.

### Transduction of human DM1 myoblasts and isolated *HSA^LR^* muscle fibers

Virus carrying dSaCas9 and control or (CAG)_6_ gRNA was used to infect human DM1 primary myoblast cell lines and *HSA*^LR^ mouse EDL muscle fibers. To determine whether blocking expression of the CTG repeats in the human myoblasts affected the presence of RNA foci and splicing of MBNL targets, cells were grown to 60% confluency on CC2 chamber slides and infected for 6 days (3 days in growth media plus 3 days in differentiation media) with viral titers of 10^9^. To analyze the effects on RNA foci in *HSA*^LR^ muscle fibers, 10-20 muscle fibers were cultured in wells of a 96 well plate and infected with a 10^9^ viral titer for 48h.

### Quantitation of Signal Intensity of Nuclear Foci in Mouse Muscle Fibers

Python scripts were written to quantitate intensity of FISH signal from RNA foci within nuclei of muscle fibers (python functions are listed below, and also see **Fig. S5**). Regions of interest were defined across each tile-scanned z-stack image obtained by confocal imaging. To identify and segment nuclei, 16-bit intensity values were scaled to lie between 0 and 1 (skimage.img_as_float). Then, a gaussian filter of standard deviation 3 pixels was applied to the raw DAPI signal (skimage.filters.gaussian), and a threshold of mean + 5 * standard deviation of the gaussian filtered signal was used to generate a binary mask. Holes were filled (scipy.ndimage.fill_holes), and a binary opening operation was performed to remove salt noise (skimage.morphology.binary_opening with ball structured element of radius 2). Nuclei were segmented and labeled (skimage.measure.label), and objects with pixel volume <1000 or >20000 were removed. FISH signal was scaled to lie between 0 and 1 as above, and a grayscale openings operation was performed to measure background (skimage.morphology.opening with a ball structured element of radius 3). This background intensity multipled by 3 was subtracted from the FISH signal to yield background-subtracted FISH signal. The binary nucleus mask was applied to this signal, and total intensity was measured within each nucleus, and divided by the nuclear volume in pixels, to obtain the final FISH signal per unit volume for each nucleus. This procedure was applied to all regions of interest across all fibers.

### Analysis of RNAseq data

100 ng of RNA was used to prepare RNA-Seq libraries using the KAPA Ribo-Erase Strand-Specific kit. Samples were pooled and sequenced on the NextSeq 500 Version 2, using a High-Output 2x75 kit. Reads were mapped to hg19 by Hisat2, and splicing events were quantitated by MISO. Ψ values from DM tibialis biopsies were fit to sigmoid curves using 4-parameter estimation, where Ψ = Ψ_min_ + (Ψ_max_ - Ψ_min_) / (1 + e^-slope^ ^*^ ^([MBNL]inferred-EC50)^), using python/scipy packages. The [MBNL]_inferred_ value was taken from Wagner et al^3434^. The “fit error” was evaluated by taking the sum of squared errors between observed Ψ and Ψ as predicted by the sigmoid curves. Events consistently regulated between non-DM1 and DM1 myoblasts were identified using a modified monotonicity test^5454^, ΔΨ > 0.1, BF > 5) where the 2 non-DM1 libraries were grouped together, and 4 DM1 libraries were grouped together. For Figure 5B, events with <1.3 sigmoid fit error and >1 monotonicity Z-score were selected for display. For Figure 5F, only events identified in Figure 5B to lie in the upper right or lower left quadrants were further analyzed; in addition, events were required to exhibit <0.33 difference in Ψ between cells treated with AAV-dSaCas9-control gRNA and non-DM1 tibialis biopsies. Raw RNA-Seq reads for these libraries are publicly available (GEO accession number pending). Statistics on read coverage are in Table S2.

### Electromyography

To determine whether expression of dCas9-(CAG)_6_ rescued myotonia in the *HSA*^*LR*^ mice, mice were injected with AAV6-dSaCas9 and control or (CAG)_6_ gRNA at 10^10^ viral genomes per mouse via the temporal vein at P2^55^. Myotonia was assessed by electromyography at 5 weeks of age. Electomyography was performed under general anaesthesia (intraperitoneal ketamine, 100 mg/kg; xylazine, 10 mg/kg) using 30 gauge concentric needle electrodes to examine the hindlimb gastrocnemius and tibialis anterior muscles. At least 15 needle insertions were performed in each muscle and myotonic discharges were denoted as a percentage of the total number of insertions.

## Acknowledgements

We would like to thank L. Ranum, T. Zu and J. Cleary for generously providing us with the plasmids to assess RAN translation, G. Aslanidi for providing the AAV6 capsid plasmid and J. Singh for advice on AAV production. We thank T. Wang for construction of the CCTG_240_ repeat tract. We thank C. Thornton for RNA samples from DM1 patients, which were obtained from the BioBank of the University of Rochester Wellstone Center (NIH NS048843). E.T.W. and T.S. were supported by DP5 OD017865. L. D. was supported by a Thomas H. Maren Fellowship.

## Author Contributions

Conceptualization, T.S., B.P., E.T.W.; Methodology, T.S., B.P., E.T.W.; Software, T.S., E.T.W.; Formal Analysis, T.S., B.P., E.T.W.; Investigation, T.S., B.P., R.O., J.C., H.M., L.D., O.M., J.O., E.T.W.; Initiated Project, T.S., E.T.W.; Resources, H.M., G.X., R.O., M.S.; Writing - Original Draft, T.S., B.P., E.T.W.; Writing - Review & Editing, T.S., B.P., E.T.W., M.S., H.M., G.X., R.O.; Visualization, T.S., B.P., E.T.W.; Funding Acquisition, E.T.W.; Supervision, E.T.W.

## References

1. Nelson, D. L., Orr, H. T., & Warren, S. T. (2013) The unstable repeats—three evolving faces of neurological disease. Neuron 77, 825–43.

2. Kennedy, L., Evans, E., Chen, C.-M., Craven, L., Detloff, P. J., Ennis, M., & Shelbourne, P. F. (2003) Dramatic tissue-specific mutation length increases are an early molecular event in Huntington disease pathogenesis. Hum Mol Genet 12, 3359–67.

3. Thornton, C. A., Johnson, K., & Moxley, 3rd, R. T. (1994) Myotonic dystrophy patients have larger CTG expansions in skeletal muscle than in leukocytes. Ann Neurol 35, 104–7.

4. Miller, J. W., Urbinati, C. R., Teng-Umnuay, P., Stenberg, M. G., Byrne, B. J., Thornton, C. A., & Swanson, M. S. (2000) Recruitment of human muscleblind proteins to (CUG)(n) expansions associated with myotonic dystrophy. EMBO J 19, 4439–48.

5. Kanadia, R. N., Johnstone, K. A., Mankodi, A., Lungu, C., Thornton, C. A., Esson, D., Timmers, A. M., Hauswirth, W. W., & Swanson, M. S. (2003) A muscleblind knockout model for myotonic dystrophy. Science 302, 1978–80.

6. Du, H., Cline, M. S., Osborne, R. J., Tuttle, D. L., Clark, T. A., Donohue, J. P., Hall, M. P., Shiue, L., Swanson, M. S., Thornton, C. A., & Ares, Jr, M. (2010) Aberrant alternative splicing and extracellular matrix gene expression in mouse models of myotonic dystrophy. Nat Struct Mol Biol 17, 187–93.

7. Wang, E. T., Cody, N. A. L., Jog, S., Biancolella, M., Wang, T. T., Treacy, D. J., Luo, S., Schroth, G. P., Housman, D. E., Reddy, S., Lécuyer, E., & Burge, C. B. (2012) Transcriptome-wide regulation of pre-mRNA splicing and mRNA localization by muscleblind proteins. Cell 150, 710–24.

8. Gendron, T. F., Belzil, V. V., Zhang, Y.-J., & Petrucelli, L. (2014) Mechanisms of toxicity in C9FTLD/ALS. Acta Neuropathol 127, 359–76.

9. Zu, T., Gibbens, B., Doty, N. S., Gomes-Pereira, M., Huguet, A., Stone, M. D., Margolis, J., Peterson, M., Markowski, T. W., Ingram, M. A. C., Nan, Z., Forster, C., Low, W. C., Schoser, B., Somia, N. V., Clark, H. B., Schmechel, S., Bitterman, P. B., Gourdon, G., Swanson, M. S., Moseley, M., & Ranum, L. P. W. (2011) Non-ATG-initiated translation directed by microsatellite expansions. Proc Natl Acad Sci U S A 108, 260–5.

10. Zu, T., Liu, Y., Bañez-Coronel, M., Reid, T., Pletnikova, O., Lewis, J., Miller, T. M., Harms, M. B., Falchook, A. E., Subramony, S. H., Ostrow, L. W., Rothstein, J. D., Troncoso, J. C., & Ranum, L. P. W. (2013) RAN proteins and RNA foci from antisense transcripts in C9ORF72 ALS and frontotemporal dementia. Proc Natl Acad Sci U S A 110, E4968–77.

11. Wheeler, T. M., Leger, A. J., Pandey, S. K., MacLeod, A. R., Nakamori, M., Cheng, S. H., Wentworth, B. M., Bennett, C. F., & Thornton, C. A. (2012) Targeting nuclear RNA for in vivo correction of myotonic dystrophy. Nature 488, 111–5.

12. Lagier-Tourenne, C., Baughn, M., Rigo, F., Sun, S., Liu, P., Li, H.-R., Jiang, J., Watt, A. T., Chun, S., Katz, M., Qiu, J., Sun, Y., Ling, S.-C., Zhu, Q., Polymenidou, M., Drenner, K., Artates, J. W., McAlonis-Downes, M., Markmiller, S., Hutt, K. R., Pizzo, D. P., Cady, J., Harms, M. B., Baloh, R. H., Vandenberg, S. R., Yeo, G. W., Fu, X.-D., Bennett, C. F., Cleveland, D. W., & Ravits, J. (2013) Targeted degradation of sense and antisense C9orf72 RNA foci as therapy for ALS and frontotemporal degeneration. Proc Natl Acad Sci U S A 110, E4530–9.

13. Sah, D. W. Y. & Aronin, N. (2011) Oligonucleotide therapeutic approaches for Huntington disease. J Clin Invest 121, 500–7.

14. Furling, D., Doucet, G., Langlois, M.-A., Timchenko, L., Belanger, E., Cossette, L., & Puymirat, J. (2003) Viral vector producing antisense RNA restores myotonic dystrophy myoblast functions. Gene Ther 10, 795–802.

15. François, V., Klein, A. F., Beley, C., Jollet, A., Lemercier, C., Garcia, L., & Furling, D. (2011) Selective silencing of mutated mRNAs in DM1 by using modified hU7-snRNAs. Nat Struct Mol Biol 18, 85–7.

16. Langlois, M.-A., Lee, N. S., Rossi, J. J., & Puymirat, J. (2003) Hammerhead ribozyme-mediated destruction of nuclear foci in myotonic dystrophy myoblasts. Mol Ther 7, 670–80.

17. Harper, S. Q., Staber, P. D., He, X., Eliason, S. L., Martins, I. H., Mao, Q., Yang, L., Kotin, R. M., Paulson, H. L., & Davidson, B. L. (2005) RNA interference improves motor and neuropathological abnormalities in a Huntington’s disease mouse model. Proc Natl Acad Sci U S A 102, 5820–5.

18. Siboni, R. B., Nakamori, M., Wagner, S. D., Struck, A. J., Coonrod, L. A., Harriott, S. A., Cass, D. M., Tanner, M. K., & Berglund, J. A. (2015) Actinomycin D Specifically Reduces Expanded CUG Repeat RNA in Myotonic Dystrophy Models. Cell Rep 13, 2386–94.

19. Rzuczek, S. G., Colgan, L. A., Nakai, Y., Cameron, M. D., Furling, D., Yasuda, R., & Disney, M. D. (2017) Precise small-molecule recognition of a toxic CUG RNA repeat expansion. Nat Chem Biol 13, 188–193.

20. Liu, C.-R., Chang, C.-R., Chern, Y., Wang, T.-H., Hsieh, W.-C., Shen, W.-C., Chang, C.-Y., Chu, I.-C., Deng, N., Cohen, S. N., & Cheng, T.-H. (2012) Spt4 is selectively required for transcription of extended trinucleotide repeats. Cell 148, 690–701.

21. Kramer, N. J., Carlomagno, Y., Zhang, Y.-J., Almeida, S., Cook, C. N., Gendron, T. F., Prudencio, M., Van Blitterswijk, M., Belzil, V., Couthouis, J., Paul, 3rd, J. W., Goodman, L. D., Daughrity, L., Chew, J., Garrett, A., Pregent, L., Jansen-West, K., Tabassian, L. J., Rademakers, R., Boylan, K., Graff-Radford, N. R., Josephs, K. A., Parisi, J. E., Knopman, D. S., Petersen, R. C., Boeve, B. F., Deng, N., Feng, Y., Cheng, T.-H., Dickson, D. W., Cohen, S. N., Bonini, N. M., Link, C. D., Gao, F.-B., Petrucelli, L., & Gitler, A. D. (2016) Spt4 selectively regulates the expression of C9orf72 sense and antisense mutant transcripts. Science 353, 708–12.

22. Qi, L. S., Larson, M. H., Gilbert, L. A., Doudna, J. A., Weissman, J. S., Arkin, A. P., & Lim, W. A. (2013) Repurposing CRISPR as an RNA-guided platform for sequence-specific control of gene expression. Cell 152, 1173–83.

23. Gilbert, L. A., Larson, M. H., Morsut, L., Liu, Z., Brar, G. A., Torres, S. E., Stern-Ginossar, N., Brandman, O., Whitehead, E. H., Doudna, J. A., Lim, W. A., Weissman, J. S., & Qi, L. S. (2013) CRISPR-mediated modular RNA-guided regulation of transcription in eukaryotes. Cell 154, 442–51.

24. Lawhorn, I. E. B., Ferreira, J. P., & Wang, C. L. (2014) Evaluation of sgRNA target sites for CRISPR-mediated repression of TP53. PLoS One 9, e113232.

25. Konermann, S., Brigham, M. D., Trevino, A. E., Hsu, P. D., Heidenreich, M., Cong, L., Platt, R. J., Scott, D. A., Church, G. M., & Zhang, F. (2013) Optical control of mammalian endogenous transcription and epigenetic states. Nature 500, 472–6.

26. Gilbert, L. A., Horlbeck, M. A., Adamson, B., Villalta, J. E., Chen, Y., Whitehead, E. H., Guimaraes, C., Panning, B., Ploegh, H. L., Bassik, M. C., Qi, L. S., Kampmann, M., & Weissman, J. S. (2014) Genome-Scale CRISPR-Mediated Control of Gene Repression and Activation. Cell 159, 647–61.

27. Chen, B., Gilbert, L. A., Cimini, B. A., Schnitzbauer, J., Zhang, W., Li, G.-W., Park, J., Blackburn, E. H., Weissman, J. S., Qi, L. S., & Huang, B. (2013) Dynamic imaging of genomic loci in living human cells by an optimized CRISPR/Cas system. Cell 155, 1479–91.

28. Kuscu, C., Arslan, S., Singh, R., Thorpe, J., & Adli, M. (2014) Genome-wide analysis reveals characteristics of off-target sites bound by the Cas9 endonuclease. Nat Biotechnol 32, 677–83.

29. Ho, T. H., Savkur, R. S., Poulos, M. G., Mancini, M. A., Swanson, M. S., & Cooper, T. A. (2005) Colocalization of muscleblind with RNA foci is separable from mis-regulation of alternative splicing in myotonic dystrophy. J Cell Sci 118, 2923–33.

30. Ho, T. H., Charlet-B, N., Poulos, M. G., Singh, G., Swanson, M. S., & Cooper, T. A. (2004) Muscleblind proteins regulate alternative splicing. EMBO J 23, 3103–12.

31. Charlet-B, N., Savkur, R. S., Singh, G., Philips, A. V., Grice, E. A., & Cooper, T. A. (2002) Loss of the muscle-specific chloride channel in type 1 myotonic dystrophy due to misregulated alternative splicing. Mol Cell 10, 45–53.

32. Savkur, R. S., Philips, A. V., & Cooper, T. A. (2001) Aberrant regulation of insulin receptor alternative splicing is associated with insulin resistance in myotonic dystrophy. Nat Genet 29, 40–7.

33. Parkesh, R., Childs-Disney, J. L., Nakamori, M., Kumar, A., Wang, E., Wang, T., Hoskins, J., Tran, T., Housman, D., Thornton, C. A., & Disney, M. D. (2012) Design of a bioactive small molecule that targets the myotonic dystrophy type 1 RNA via an RNA motif-ligand database and chemical similarity searching. J Am Chem Soc 134, 4731–42.

34. Wagner, S. D., Struck, A. J., Gupta, R., Farnsworth, D. R., Mahady, A. E., Eichinger, K., Thornton, C. A., Wang, E. T., & Berglund, J. A. (2016) Dose-Dependent Regulation of Alternative Splicing by MBNL Proteins Reveals Biomarkers for Myotonic Dystrophy. PLoS Genet 12, e1006316.

35. Ran, F. A., Cong, L., Yan, W. X., Scott, D. A., Gootenberg, J. S., Kriz, A. J., Zetsche, B., Shalem, O., Wu, X., Makarova, K. S., Koonin, E. V., Sharp, P. A., & Zhang, F. (2015) In vivo genome editing using Staphylococcus aureus Cas9. Nature 520, 186–91.

36. Nishimasu, H., Cong, L., Yan, W. X., Ran, F. A., Zetsche, B., Li, Y., Kurabayashi, A., Ishitani, R., Zhang, F., & Nureki, O. (2015) Crystal Structure of Staphylococcus aureus Cas9. Cell 162, 1113–26.

37. Mankodi, A., Logigian, E., Callahan, L., McClain, C., White, R., Henderson, D., Krym, M., & Thornton, C. A. (2000) Myotonic dystrophy in transgenic mice expressing an expanded CUG repeat. Science 289, 1769–73.

38. Chew, W. L., Tabebordbar, M., Cheng, J. K. W., Mali, P., Wu, E. Y., Ng, A. H. M., Zhu, K., Wagers, A. J., & Church, G. M. (2016) A multifunctional AAV-CRISPR-Cas9 and its host response. Nat Methods 13, 868–74.

39. Duddy, W., Duguez, S., Johnston, H., Cohen, T. V., Phadke, A., Gordish-Dressman, H., Nagaraju, K., Gnocchi, V., Low, S., & Partridge, T. (2015) Muscular dystrophy in the mdx mouse is a severe myopathy compounded by hypotrophy, hypertrophy and hyperplasia. Skelet Muscle 5, 16.

40. van Agtmaal, E. L., André, L. M., Willemse, M., Cumming, S. A., van Kessel, I. D. G., van den Broek, W. J. A. A., Gourdon, G., Furling, D., Mouly, V., Monckton, D. G., Wansink, D. G., & Wieringa, B. (2017) CRISPR/Cas9-Induced (CTG·CAG)n Repeat Instability in the Myotonic Dystrophy Type 1 Locus: Implications for Therapeutic Genome Editing. Mol Ther 25, 24–43.

41. Cinesi, C., Aeschbach, L., Yang, B., & Dion, V. (2016) Contracting CAG/CTG repeats using the CRISPR-Cas9 nickase. Nat Commun 7, 13272.

42. Batra, R., Nelles, D. A., Pirie, E., Blue, S. M., Marina, R. J., Wang, H., Chaim, I. A., Thomas, J. D., Zhang, N., Nguyen, V., Aigner, S., Markmiller, S., Xia, G., Corbett, K. D., Swanson, M. S., & Yeo, G. W. (2017) Elimination of Toxic Microsatellite Repeat Expansion RNA by RNA-Targeting Cas9. Cell 170, 899–912.e10.

43. Kemaladewi, D. U., Maino, E., Hyatt, E., Hou, H., Ding, M., Place, K. M., Zhu, X., Bassi, P., Baghestani, Z., Deshwar, A. G., Merico, D., Xiong, H. Y., Frey, B. J., Wilson, M. D., Ivakine, E. A., & Cohn, R. D. (2017) Correction of a splicing defect in a mouse model of congenital muscular dystrophy type 1A using a homology-directed-repair-independent mechanism. Nat Med 23, 984–989.

44. Tabebordbar, M., Zhu, K., Cheng, J. K. W., Chew, W. L., Widrick, J. J., Yan, W. X., Maesner, C., Wu, E. Y., Xiao, R., Ran, F. A., Cong, L., Zhang, F., Vandenberghe, L. H., Church, G. M., & Wagers, A. J. (2016) In vivo gene editing in dystrophic mouse muscle and muscle stem cells. Science 351, 407–411.

45. Xia, G., Santostefano, K. E., Goodwin, M., Liu, J., Subramony, S. H., Swanson, M. S., Terada, N., & Ashizawa, T. (2013) Generation of neural cells from DM1 induced pluripotent stem cells as cellular model for the study of central nervous system neuropathogenesis. Cell Reprogram 15, 166–77.

46. Philips, A. V., Timchenko, L. T., & Cooper, T. A. (1998) Disruption of splicing regulated by a CUG-binding protein in myotonic dystrophy. Science 280, 737–41.

47. Mali, P., Yang, L., Esvelt, K. M., Aach, J., Guell, M., DiCarlo, J. E., Norville, J. E., & Church, G. M. (2013) RNA-guided human genome engineering via Cas9. Science 339, 823–6.

48. Cheng, A. W., Wang, H., Yang, H., Shi, L., Katz, Y., Theunissen, T. W., Rangarajan, S., Shivalila, C. S., Dadon, D. B., & Jaenisch, R. (2013) Multiplexed activation of endogenous genes by CRISPR-on, an RNA-guided transcriptional activator system. Cell Res 23, 1163–71.

49. Rutledge, E. A., Halbert, C. L., & Russell, D. W. (1998) Infectious clones and vectors derived from adeno-associated virus (AAV) serotypes other than AAV type 2. J Virol 72, 309–19.

50. Xiao, X., Li, J., & Samulski, R. J. (1998) Production of high-titer recombinant adeno-associated virus vectors in the absence of helper adenovirus. J Virol 72, 2224–32.

51. Zolotukhin, S., Potter, M., Zolotukhin, I., Sakai, Y., Loiler, S., Fraites, Jr, T. J., Chiodo, V. A., Phillipsberg, T., Muzyczka, N., Hauswirth, W. W., Flotte, T. R., Byrne, B. J., & Snyder, R. O. (2002) Production and purification of serotype 1, 2, and 5 recombinant adeno-associated viral vectors. Methods 28, 158–67.

52. Piedra, J., Ontiveros, M., Miravet, S., Penalva, C., Monfar, M., & Chillon, M. (2015) Development of a rapid, robust, and universal picogreen-based method to titer adeno-associated vectors. Hum Gene Ther Methods 26, 35–42.

53. Pasut, A., Jones, A. E., & Rudnicki, M. A. (2013) Isolation and culture of individual myofibers and their satellite cells from adult skeletal muscle. J Vis Exp, e50074.

54. Wang, E. T., Ward, A. J., Cherone, J. M., Giudice, J., Wang, T. T., Treacy, D. J., Lambert, N. J., Freese, P., Saxena, T., Cooper, T. A., & Burge, C. B. (2015) Antagonistic regulation of mRNA expression and splicing by CELF and MBNL proteins. Genome Res 25, 858–71.

55. Gombash Lampe, S. E., Kaspar, B. K., & Foust, K. D. (2014) Intravenous injections in neonatal mice. J Vis Exp, e52037.

